# Mechanism of branching morphogenesis inspired by diatom silica formation

**DOI:** 10.1101/2023.07.03.547407

**Authors:** Iaroslav Babenko, Nils Kröger, Benjamin M. Friedrich

**Author notes:** Corresponding authors: Nils Kröger, Benjamin M. Friedrich.

## Abstract

The silica-based cell walls of diatoms are prime examples of genetically controlled, species-specific mineral architectures. The physical principles underlying morphogenesis of their hierarchically structured silica patterns are not understood, yet such insight could indicate novel routes towards synthesizing functional inorganic materials. Recent advances in imaging nascent diatom silica allow rationalizing possible mechanisms of their pattern formation. Here, we combine theory and experiments on the model diatom *Thalassiosira pseudonana* to put forward a minimal model of branched rib patterns – a fundamental feature of the silica cell wall. We quantitatively recapitulate the time-course of rib pattern morphogenesis by accounting for silica biochemistry with autocatalytic formation of diffusible silica precursors followed by conversion into solid silica. We propose that silica deposition releases an inhibitor that slows down up-stream precursor conversion, thereby implementing a self-replicating reaction-diffusion system featuring a non-classical Turing mechanism. The proposed mechanism highlights the role of geometrical cues for guided self-organization, rationalizing the instructive role for the single initial pattern seed known as primary silicification site. The mechanism of branching morphogenesis that we characterize here is possibly generic and may apply also in other biological systems.

**Significance statement:** The formation of minerals by living organisms is a widespread biological phenomenon occurring throughout the evolutionary tree-of-life. The silica-based cell walls of diatom microalgae are impressive examples featuring intricate architectures and outstanding materials properties that still defy their reconstitution *in vitro*. Here, we developed a minimal mathematical model that explains the formation of branched patterns of silica ribs, providing unprecedented understanding of basic physico-chemical processes capable of guiding silica morphogenesis in diatoms. The generic mechanism of branching morphogenesis identified here could provide recipes for bottom-up synthesis of mineral-nanowire networks for technological applications. Moreover, similar mechanisms may apply in the biological morphogenesis of other branched structures, like corals, bacterial colonies, tracheal networks, fungal plexuses, or the vascular system.

## Introduction

Diatoms are unicellular algae that produce intricately patterned cell walls made predominantly of amorphous silica (SiO_2_). These photosynthetic microorganisms can be found in all aquatic habitats and play an important ecological role, with marine diatoms alone being responsible for about 20% of marine carbon fixation [1], [2]. The architectural intricacy of diatom silica is unique, yet its biological functions have remained largely speculative. The cell wall serves likely as protective armor against zooplankton predators [3], [4] and might also play a role in nutrient uptake [5], enhance light harvesting for photosynthesis [6] and protect the cell from harmful ultraviolet radiation [7]. Due to its unique material properties, synthetically modified diatom silica is being explored for advanced technological applications including biomedicine [8], [9].

Diatoms represent a diverse group with estimated 30,000 species [10], [11], each producing a specific cell wall morphology with characteristic patterns of nano- and microscale features. Despite the large structural variety, the basic cell wall architecture is conserved, consisting of two interlocking halves, each comprising a valve and several connecting girdle bands (Fig. 1A). While girdle bands commonly possess a plain morphology with smooth surfaces and uniform patterns of pores, valves are intricately patterned, typically with a network of silica ribs and intervening hierarchical arrangements of meso- and nanoscale pores [12]–[14]. Valves and girdle bands are produced in separate intracellular compartments each termed silica deposition vesicle (SDV) [12]. The valve SDV is disk-shaped, and expands concurrently with the nascent silica patterns that develop therein (Fig. 1B). It has been proposed that transport vesicles delivering silicic acid continuously fuse with the SDV, thus driving its expansion [15]. Once valve morphogenesis inside the SDV is complete, the new silica valve is exocytosed and integrated into the cell wall [12], [13]. In recent years, tremendous progress has been made in diatom research regarding sequencing and targeted manipulation of genomes [16], as well as the identification of candidate silica biogenesis proteins [17]–[19]. However, the physical principles that govern morphogenesis of the hierarchical silica patterns in diatoms are still enigmatic.

**Figure 1:**
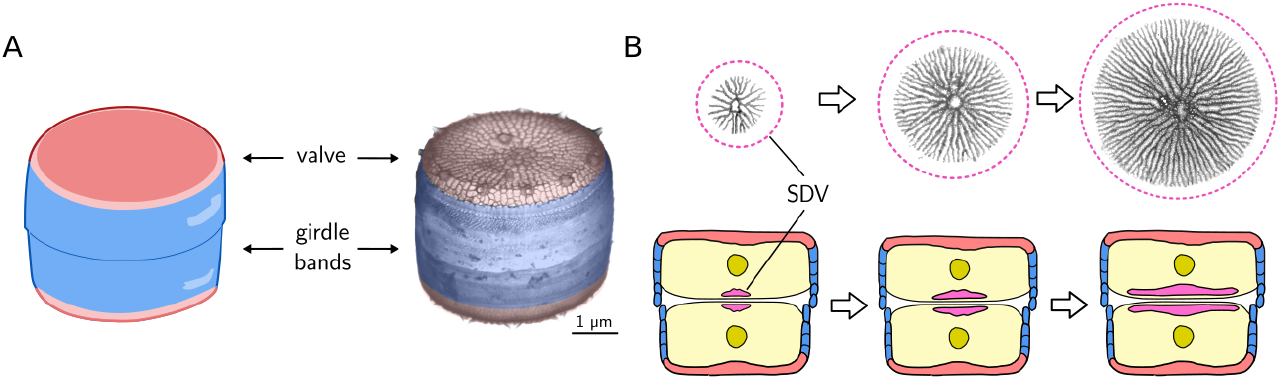
Diatom cell wall morphogenesis. (A) Schematic of the cell wall (left) and scanning electron microscopy (SEM) image (right) of *T. pseudonana*, highlighting valves (red) and girdle bands (blue). (B, top) TEM images of valve SDVs showing nascent silica rib networks. The putative outline of the SDV membrane is indicated in purple. (B, bottom) Schematic showing a diatom cell in cross-section shortly after cytokinesis, highlighting the expanding valve SDVs in each daughter protoplast.

Previous attempts to model valve morphogenesis focused on replicating the patterns in the final, mature valve rather than trying to mimic the temporal dynamics of its formation. Early diffusion-limited aggregation models produced irregular, fractal-like rib patterns, which do not yet match the regular rib patterns of diatom valves [20], [21]. Moreover, these models assumed silica influx from the rim of the SDV, for which there is no experimental support. Bentley and co-workers addressed cell-scale patterns using a cellular automaton model, but had to impose pre-patterns of dissolvable organic material to direct the formation of so-called virgae ribs [22]. Other studies investigated phase-separation as a mechanism for valve morphogenesis addressing formation of pores [23], [24] but did not account for branching patterns of silica. Others put their efforts exclusively on studying the formation of regular pore patterns assuming a pre-pattern of silica ribs [23], [25], [26].

Recently, a method was developed that enabled imaging silica morphogenesis in the valve SDVs of the model diatom *Thalassiosira pseudonana* at various developmental stages in unprecedented detail using transmission electron microscopy (TEM) [27]. This provides a pseudotime series of silica formation, which allows rationalizing physico-chemical mechanisms of valve morphogenesis. As is the case in all diatoms species with radial valve symmetry (centric diatoms), valve morphogenesis starts from a central, circular seed and expands as a network of branching silica ribs with constant spacing [27]–[30]. The initial seed is known as the primary silicification site (PSS) and obscured in the mature valve (Fig. 1A).

Here, we aimed to establish a minimal mathematical model that – based on known silica biochemistry – quantitatively accounts for the pseudo-time course of the experimentally observed nascent silica rib pattern in *T. pseudonana* valve SDVs. Such a model may reveal general physical principles of silica morphogenesis in diatoms.

## Results

### Model of branching morphogenesis

One of the most conspicuous features of nascent valve silica is the regular spacing between adjacent ribs. Spontaneous formation of regularly spaced structures is reminiscent of Turing patterns, which originate from reaction-diffusion systems with nonlinear feedback [31]–[33]. It is tempting to assume that silica deposition in diatoms is governed by a similar mechanism. However, classical Turing systems predict the formation of multiple seeds from slightly inhomogeneous initial conditions, whereas the silica rib patterns in diatom valves always start from a single initial seed. Therefore, self-replicating reaction-diffusion systems that require a strong local perturbation (i.e. “non-classical” Turing systems) [34]–[36] provide a more suitable approach to rationalize diatom valve morphogenesis. We propose a putative mechanism for branching morphogenesis of silica rib patterns in terms of a minimal mathematical model supported by experimental data (Fig. 2A) and known silica biochemistry (reviewed below). We assume that a precursor for solid silica is continuously delivered to the morphogenetic compartment as a soluble, diffusible substance 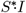, with a rate that keeps the concentration of the precursor constant within the compartment. Inside the compartment, the prevalent physico-chemical conditions cause the precursor to transition into a metastable, diffusible component 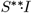 that is prone to solidify. Upon subsequent conversion of this metastable component into solid silica, an inhibitor *I* is released that slows downs the precursor conversion.

**Figure 2:**
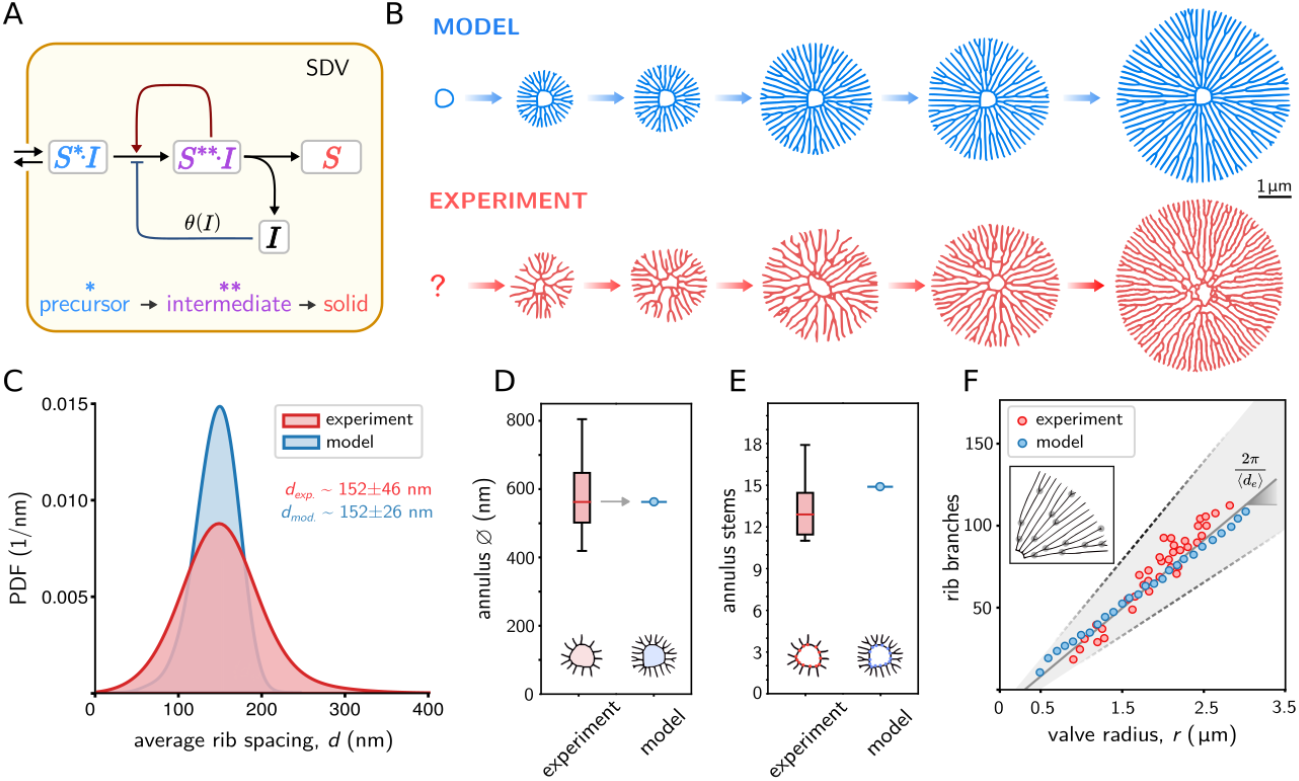
Minimal model of rib pattern morphogenesis. (A) Proposed reaction scheme, comprising silica precursor 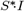, soluble silica 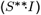, and solidified silica (*S*). Inside the SDV a constant precursor concentration is converted into solid silica, thereby releasing an inhibitor (*I*) that slows down precursor conversion. (B) Simulated rib patterns using Eq. 0.1 (blue), together with pseudotime course of nascent rib networks from skeletonized TEM images representing subsequent stages of valve development (red). (C) Spacing between neighboring ribs in experimentally observed rib patterns (red, n=38 valves) pooled over different developmental stages, and in simulated patterns (blue, n=28). The mean spacing was used to calibrate unit length in simulations. (D) Diameter of the central annulus observed in experiments (red, n=23), and simulations (blue). The annulus diameter was used to scale initial conditions in simulations (gray arrow). (E) Number of initial stems of the central annulus in experiments (red, n=23) and simulations. (F) Number of rib branches *N*_*b*_ (stem branching points excluded) as a function of valve radius *r* for experimentally observed patterns (red), simulated patterns (blue), as well as the prediction from a simple geometric law (black).

In the scope of reaction-diffusion systems, a linear reaction chain 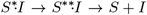 cannot form patterns without inherent autocatalytic feedback. Yet, a simple nonlinear feedback in the formation of the metastable component provides a reaction-diffusion system capable of self-organized pattern formation. We postulate that the metastable component 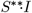 accelerates its own production via autocatalytic activity which can be inhibited by free inhibitor *I*.

The model is compatible with known silica biochemistry in diatoms, where the SDV represents the morphogenetic compartment. Complexes of the inhibitor with monosilicic acid or low molecular mass silicic acid molecules could represent the silica precursor 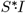. Such complexes would be stable in the near neutral pH of the cellular cytoplasm constituting the previously described soluble silicon pools of diatom cells [37]–[39], but become metastable upon entering the acidic lumen of the SDV [40], [41]. Inside the SDV, the precursor would undergo auto-polycondensation reactions generating a diverse pool of soluble oligo- and poly-silicic acid molecules representing the metastable intermediate 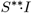. The intermediates would still retain the inhibitor molecules albeit increasingly low affinity as the degree of polycondensation increases. Finally, high molecular mass poly-silicic acid molecules aggregate and become deposited as dense, insoluble silica while releasing the inhibitor. Intriguingly, oligo- and polysilicic acid possess numerous hydroxyl groups, whose reactivity towards polycondensation increases with increasing degree of polymerization [42, p. 214]. This provides a rationale for the postulated autocatalytic self-enhancement of the conversion of silica precursor into intermediate. The inhibitor molecules could slow down oligo- and poly-silicic formation by reversibly binding to the hydroxyl groups of the silica precursor thereby enhancing its stability. The full reaction scheme of biosilica deposition is likely to be more complex, yet we anticipate that the minimal model recapitulates its key features. In particular, this model is general, posits testable hypotheses regarding positive and negative feedback, and may apply in similar form in other systems.

### Quantitative image analysis of nascent silica valves

We took advantage of a new protocol to extract valve SDVs in synchronized cell cultures of the centric diatom *T. pseudonana*, which enabled TEM imaging of nascent valves at different developmental stages [27]. We obtained a pseudotime-course of valve morphogenesis from individual TEM images by size-sorting according to valve total area *A*, exploiting the fact that valve patterns grow radially outwards.

To quantitatively characterize developing silica valves, we developed an automated image analysis algorithm to skeletonize their rib patterns (SI Appendix Fig. S1), which allowed monitoring valve morphogenesis as series of size-sorted skeletons (Fig. 2B). From this, we obtained statistics for the lateral spacing between neighboring ribs (Fig. 2C), the diameter of the central ring (annulus, Fig. 2D), the number of stems emanating from that annulus (Fig. 2E), and the number of branching points as function of valve radius *r* (Fig. 2F). We found that the rib spacing *d* is narrowly distributed, with an average of 152 nm that does not change substantially during valve development (SI Appendix, Tab. S1). The central annuli had a median diameter of 560 nm with an average of 13 initial rib stems. The number of branching points *N*_*b*_(*r*) increased linearly as function of the effective valve radius *r* (determined from valve area *A* = *πr*^2^). This observation can be rationalized as follows: assuming that the average rib spacing *d* remains constant during morphogenesis, a simple geometric argument implies that the number of ribs intersecting the circumference of a circle with a radius *r* should be 2*π rd*. Hence, there should be on average one additional branching point whenever the radius increases by Δ*r* = *d*/2*π*. This yields *N*_*b*_(*r*) = *N*_*b*_(0)+2*π r*/*d* for the expected number of branching points in a valve of area *A* = *π r*^2^, up to a constant offset *N*_*b*_(0) which corresponds to stem branching points of the annulus. A similar geometric law was previously applied to the number of defects in pore patterns of diatom valves [26].

### Mathematical description

The proposed dynamics of silica precursor, intermediate and solid silica can be cast into a system of partial differential equations

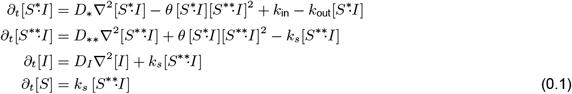

Here, the slow-down factor *θ* = *θ*([*I*]) describes the negative feedback of the inhibitor on the conversion of precursor to intermediate. We approximate its dependence on inhibitor concentration [*I*] by a sigmoid function

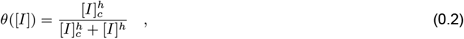

where the maximal value *θ* = 1 corresponds to unrestrained conversion of the precursor, while the minimal value *θ* = 0 corresponds to complete inhibition of precursor conversion.

### Model predictions

With a minimal set of assumptions, the model can recapitulate the temporal evolution of branched rib patterns as observed in our experiments (Fig. 2B). We used inter-rib spacing *d* and the diameter of the central annulus to calibrate model parameters (Fig. 2C,D). Interestingly, the model is capable of spontaneously forming initial stems emanating from the smooth boundary of the PSS annulus. The number of stems in simulated patterns was 16, which is comparable, but slightly larger than the mean number of rib stems observed experimentally (13*±*2.5, Fig. 2E). The number of initial rib stems in simulations was equal for perfectly circular annuli or deformed annuli of similar shape (SI Appendix, Fig. S5). Finally, we compared the number *N*_*b*_ of branching points of silica ribs as function of effective valve radius *r* for experimentally observed patterns and simulated patterns (Fig. 2F). The predicted branching frequency agreed with the measured frequency within experimental uncertainties.

The model is robust to parametric variations characterizing the inhibitor feedback, though the Gray-Scott dynamics of precursor conversion needs to be tuned to a regime generating self-replicating dot patterns (SI Appendix, Fig. S6).

For simplicity and generality, we assumed an unbounded domain in the simulations. Yet, valve morphogenesis in diatoms proceeds in an expanding compartment (i.e., the SDV). We confirmed that analogous rib patterns emerge in an expanding simulation domain with reflecting boundary conditions (SI Appendix, Fig. S7). Interestingly, the model requires continuous removal of inhibitor, which can occur either by a diffusion flux in an unbounded domain (Fig. 2), or by explicitly accounting for inhibitor degradation at a constant rate in a bounded domain (SI Appendix, Fig. S7). In fact, the removal of inhibitor may be a prerequisite for the formation of additional morphological silica features that form later, such as transverse connections between ribs and the porous silica plate filling the voids between ribs.

### Inhibitor controls rib branching

To gain insight into the mechanism of rib branching, we followed concentration profiles of all four components at the tip of a propagating rib (Fig. 3A,B). For a rib that is only propagating but not branching, the concentration profiles of the products 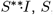, and *I* have a single peak, reflecting the width of the rib, while the precursor 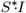 becomes locally depleted due to its conversion into intermediate. Eventually, the rib widens, causing an increase of inhibitor concentration, which slows down precursor conversion locally, causing a noticeable dent in the center of the concentration profile of the intermediate 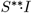. Subsequently, a dent forms also in the concentration profile of the solid product *S*. The concentration of precursor increases at the center of the rib, as less precursor is consumed. The nonlinear feedback in the reaction kinetics amplifies these effects, causing a further deepening of the dents in the concentration profiles with concomitant increase of precursor concentration between the two emerging peaks. Finally, two ribs have formed that start to repel each other.

**Figure 3:**
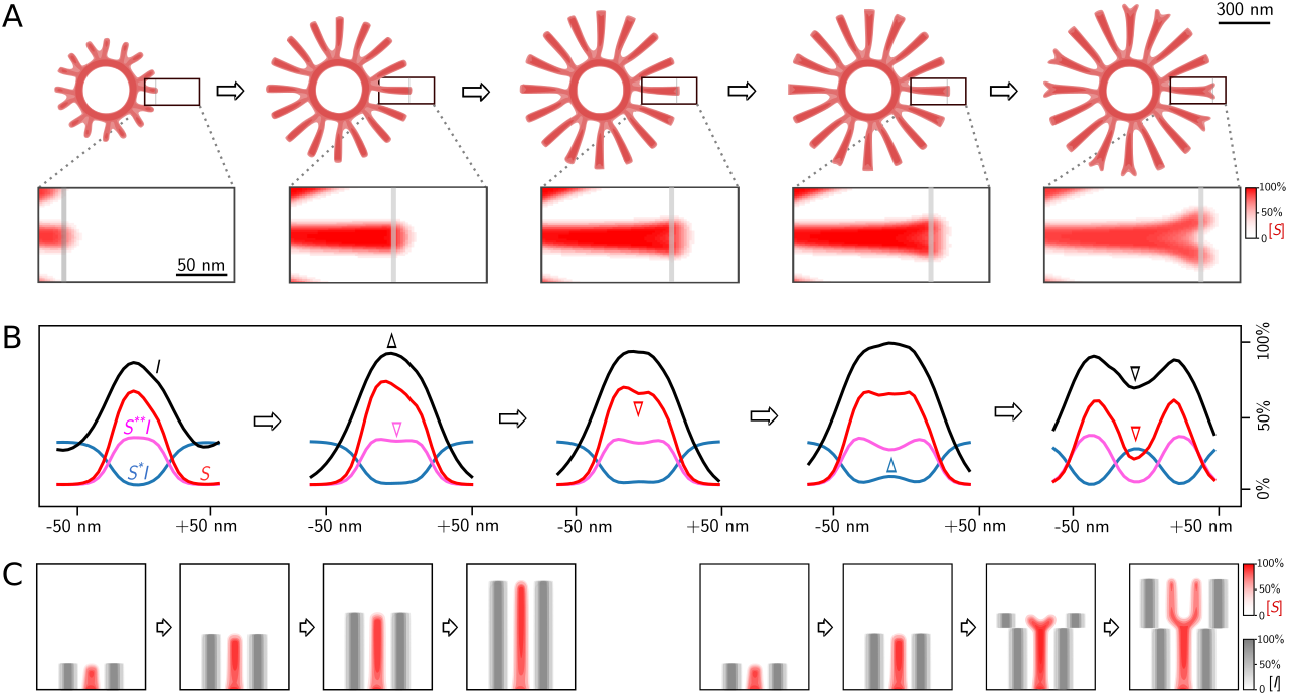
Mechanism of rib branching. (A) Simulated time-series of a branching rib pattern (showing normalized concentration of solid *S*, see color map), together with region-of-interest (ROI). (B) Normalized concentration profiles of precursor (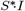, blue), intermediate (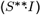, magenta), solid silica (*S*, red), and inhibitor (*I*, black) across the line-scans indicated in gray in the ROIs in panel A. Arrowheads of corresponding color highlighted changes in concentration profiles. (C) Control of rib branching by inhibitor concentration. *Left:* “artificial” ribs consisting of imposed inhibitor profiles (gray) at a distance equal to the typical inter-rib spacing from a propagating rib suppresses branching of this rib. *Right:* Increasing the lateral distance of the “artificial” ribs immediately triggers branching of the propagating rib.

In the simulated valve patterns, branching of ribs seem to occur exactly when the distance between neighboring ribs becomes larger than the typical inter-rib spacing. This implies a form of chemical communication between neighboring ribs. One potential mechanism could be a competition between neighboring ribs for precursor material. We disproved this possibility by imposing artificial sinks of precursor next to a simulated solitary rib, which virtually did not change the branching behavior (SI Appendix, Fig. S8). In contrast, imposing artificial sources of inhibitor allowed us to precisely control rib branching (Fig. 4C). Specifically, we conducted two computer experiments, where a propagating rib was flanked by ghost ribs consisting of additional inhibitor. In a first case, the distance between the ghost ribs and the propagating rib was kept constant, which reliably suppressed branching. In a second case, this distance was increased at a certain time-point, which immediately triggered branching of the central rib.

**Figure 4:**
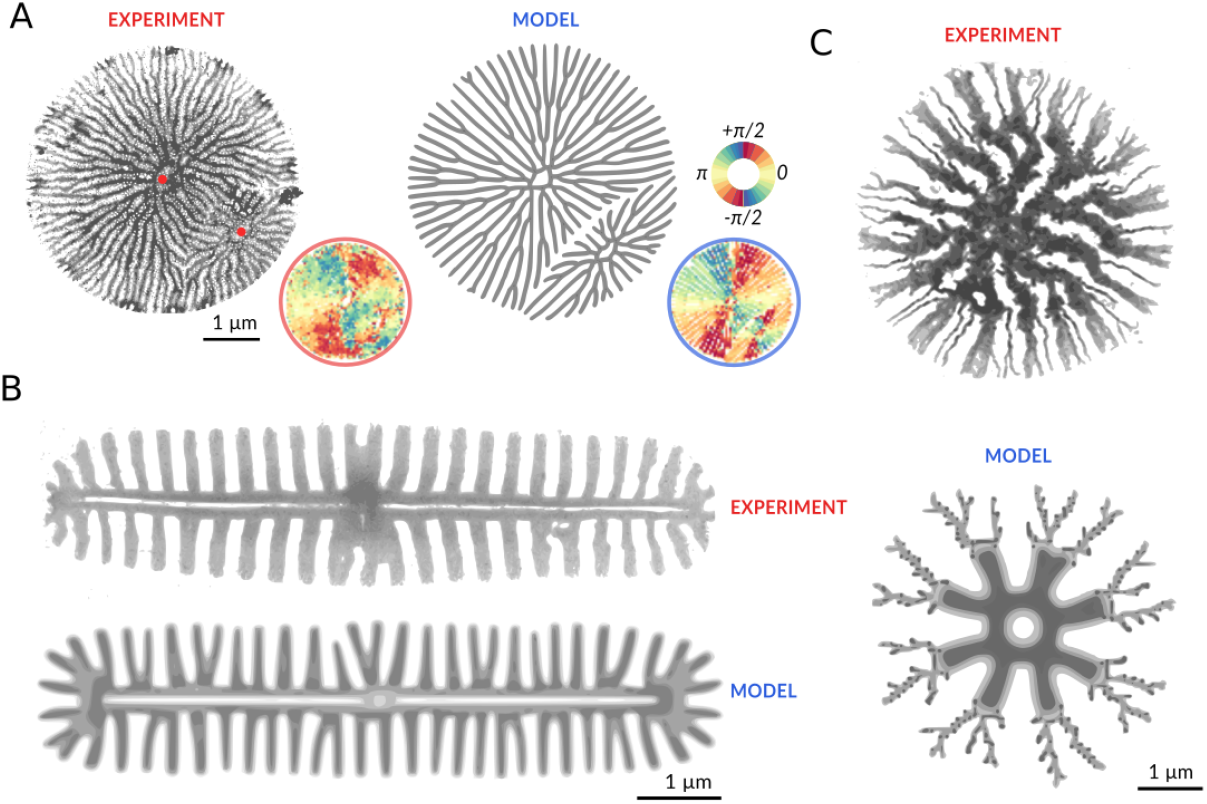
Modeling other valve silica patterns. (A, left) TEM image of an aberrant nascent *T. pseudonana* valve containing two PSSs (red dots). (A, right) simulated rib patterns in a valve using these two PSS as initial condition. The mean nematic direction for both patterns is shown for comparison below on the bottom right of each image (for reference see color wheel on the top right). (B) Silica rib pattern in the nascent valve of the pennate diatom *A. sibiricum* [28] (left), and simulated pattern using an elongated PSS with non-smooth boundary (right). (C) Silica pattern in the nascent valve of the centric diatom *C. cryptica* [27] (left), and simulated pattern using a dynamic switch of parameters (right).

### Aberrant valve morphologies and silica patterns in other diatom species

To further test the predictive capacity of the model, we investigated aberrant valve morphologies, which sporadically occur in *T. pseudonana*, as well as valve patterns in other species (Fig. 4). The example of an aberrant valve shown in Fig. 4A likely resulted from a second, ectopic PSS. Correspondingly, we simulated our model in a bounded circular domain with initial conditions given by two PSS that form shortly after another. To quantitatively compare the observed and the simulated aberrant valve pattern, we computed maps of local rib orientation, which displayed similar patterns.

The model can also partially reproduce valve patterns in other diatom species. Fig. 4B shows the silica pattern in the nascent valve of the pennate (i.e. bilaterally symmetric) diatom *Achnanthidium sibiricum*. We initialized simulations with an elongated PSS with high aspect ratio (SI Appendix, Fig. S9), which is typical for pennates [28], [43], [44]. The simulation resulted in similar patterns. Of note, as the boundary of this PSS is mostly straight, the roughness of its boundary becomes important in controlling the emergence of initial stems.

The valves of the centric diatom *Cyclotella cryptica* exhibit two types of silica ribs: narrow and wide ribs, which are regularly spaced in an alternating manner [27]. During valve development, the pattern of wide ribs is laid down first before the narrow ribs start to branch out from the wide ribs (Fig. 4C) [27]. We hypothesize that after an initial phase chemical conditions change, resulting in an abrupt switch in the width of the growing ribs. Correspondingly, we simulated the model with a change of parameters at an intermediate time point, resulting in a valve pattern that combines thick and thin ribs (Fig. 4C).

## Discussion

Here, we introduced a minimal model for morphogenesis of branched silica rib patterns, which are pervasive features of diatom cell walls. Despite its minimalistic nature, the model quantitatively recapitulates the experimentally observed rib pattern formation in the model diatom *T. pseudonana*, and is also capable of mimicking key features in other diatom species. The model makes several predictions, and supports or questions previous hypotheses, as explained in the following:

(i) It has so far been unresolved whether silica patterns of valves develop from a singular primary silicification site (PSS, [13]), or whether silica formation is initiated at multiple sites [45], [46]. Our model demands the presence of a single PSS, which orchestrates the emerging radial rib pattern. Occasionally, individual diatom cells form aberrant valves with two pattern centers. This phenomenon can be readily explained by our minimal model as the result of two competing rib patterns (see Fig. 4A).

(ii) Previously, it has been assumed that silica morphogenesis inside the valve SDV is dependent on directional influx of silica precursor [47]. Similarly, it was proposed that cytoskeletal fibers may guide pattern formation [26], [48]. While the cytoskeleton could contribute to material delivery to the SDV and determining SDV shape, our model shows that neither directed silica influx nor a direct guidance role of cytoskeletal fibers is required to explain the formation of radial rib patterns.

(iii) As explained in the results section, autocatalytic rate enhancement is an inherent chemical property of silica formation from soluble silica precursors in aqueous systems. The silicification reaction proceeds through numerous oligo- and poly-silicic acid intermediates that show increasing reactivity with the precursor [42]. Based on the following experimental data, we hypothesize that the precursor might consist of polyamine-stabilized silica nanoparticles bound to polyanionic proteins: NMR studies and subcellular fractionation of diatoms identified soluble silica nanoparticles as credible silica precursors [49], [50]; polyamines stabilize silica nanoparticles and prevent them from forming insoluble silica at neutral pH *in vitro* [51]; at SDV-like acidic pH polyanionic proteins isolated from diatom silica either accelerated or slowed down silica deposition *in vitro* depending on the protein:polyamine ratio [52], [53].

(iv) The inhibitor is a key component of our model. We anticipate that it has a dual function: it stabilizes the silica precursor at the near neutral pH outside the SDV, and, when set free during silica deposition, slows down the autocatalytic formation of the intermediate inside the SDV. The inhibitor mediates chemical communication between neighboring ribs to set a uniform rib spacing. This requires continuous removal of inhibitor, which is probably important also to allow for formation of other silica structures. Several studies speculated that phase-separated droplets may serve as templates for the pore patterns in the valve [19], [54], [55]. It is therefore possible that the inhibitor is not degraded but removed from the bulk phase by phase-separating into droplets, thus serving a third function.

(v) Our model demands that solid silica precipitates as a non-diffusible, chemically inert species.

From a broader perspective, the proposed model provides mechanistic insight into a generic mechanism of branching morphogenesis. In a nutshell, inhibitor released from neighboring ribs constrains the width of a propagating rib, which suppresses its branching. Whenever the distance to the neighbors exceeds a critical threshold, the rib broadens, causing local accumulation of inhibitor whose negative feedback drives tip splitting.

This model represents a self-replicating reaction-diffusion system [34]–[36], which we like to call non-classical Turing system, as pattern formation is only initiated in response to a sufficiently strong perturbation. This unusual feature ensures robustness to inherent concentration fluctuations (SI Appendix, Fig. S10), and provides an example of guided self-organization, where initial and boundary conditions guide pattern formation.

Reaction-diffusion models of self-organized pattern formation commonly employ either activator-inhibitor dynamics [31], or rely on substrate-depletion [32], [56]. Our model integrates both concepts, and thereby confers unprecedented control of emergent patterns, as stripe formation and branching are controlled by different chemical species.

Our minimal model is conceptually simpler than a class of reaction-diffusion models proposed previously to address branching morphogenesis [56]–[58] as it comprises a linear reaction chain with autocatalytic enhancement and a single negative feedback loop through the inhibitor. The model obeys mass balance, and makes explicit how transient dynamics builds solid-like structures.

A second popular class of branching morphogenesis models are receptor-ligand models of Schnakenberg-type with substrate depletion, used to describe lung morphogenesis and angiogenesis [32], [59] and other biological branching phenomena. Despite their formal similarity to the Gray-Scott dynamics used here for precursor and intermediate, these models exhibit a classical diffusion-driven Turing instability. Thus, in contrast to our model, pattern formation would start simultaneously in the entire domain in the presence of noise. Indeed, compartimentalization was proposed to increase the robustness of these models, with Turing instabilities driving the dynamics of interfaces [60], [61].

The chemical reaction scheme proposed in our model has the emergent property that tips start to branch whenever the distance to their neighbors exceeds a critical threshold. A number of previous models of branching morphogenesis evoked this emergent property as a heuristic rule to successfully predict network statistics [62]–[65]. It is tempting to speculate that the branching mechanisms introduced by our model may apply in other biological systems [66]–[68], with the model components assigned to different roles, e.g., different differentiation stages of cells, chemical signals, or transcription factors.

## Materials and Methods

### Imaging of nascent silica

Visualization of synchronized cell cultures and correlative fluorescence and electron microscopy of valve SDVs were performed as described previously [27].

### Image analysis of TEM images and the model

TEM images of nascent silica valves were analyzed using custom-made automatic skeleton recognition (python v3.7.4, using scikit-image package v0.15.0 [69]). First, a binary mask of the rib network was determined using adaptive thresholding and the rib skeleton determined using a pixel thinning algorithm. The obtained skeleton was then pruned to remove short branches up to 50 nm in length (Fig. S1). In total we were able to extract skeletons from 38 valves (Fig. S2). The inter-rib spacing was calculated from the inverted skeleton mask, followed by application of the median-axes function, which provided a graph of points equally distant to adjacent ribs (Fig. S3). The number of branching points, cross-connections and terminal points was computed from the skeleton mask using the mahotas python library (v1.4.11; pixel-type connectivity) [70] with custom kernels for pixel connectivity (Fig. S4).

The effective valve radius *r* = (*A*/*π*)^1/2^ was calculated from the area *A* of the convex-hull of the skeleton mask. Due to the relatively high level of silicification in the center of the valve, skeletons of the annuli were traced manually (n=23, from the total of n=38) and then analyzed analogously. For Fig. 4A, the mean local rib orientation was computed using a histogram of oriented gradients.

### Numerical integration of partial differential equations

To numerically integrate (0.1), we used a forward Euler scheme with constant time step and regular grid, with three-point stencil for the diffusion operator and reflecting boundary conditions. Integration was carried out in a large static square domain (Figs. 2,3), or in a dynamic disk-shaped domain that expands at constant speed (Fig. S7). To mitigate grid effects, small noise terms were added to diffusion coefficients as in [71], [72].

The initial seed representing the PSS was chosen as 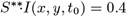 for all pixels (*x, y*) belonging to digitally-drawn, distorted annuli of thickness Δ*r* = 3, and 0 otherwise. The initial concentration of 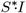 was set to stationary state 1 for the entire domain; initial concentrations were set to zero for *S* and *I*.

Parameters for Figs. 2 and 3: *k*_in_ = *k*_out_ = 0.073, *k*_sol_ = 0.135, [*I*]_*c*_ = 7.3, *h* = 55, *D*_*_ = 0.1, *D*_**_ = 0.57, *D*_*I*_ = 0.21. Numerical computations were carried out in normalized units; unit length was calibrated afterwards to match mean inter-rib spacing in Fig. 2C. Parameters used in Fig. 4 for other species are listed in SI Appendix.

### Data, Materials and Software Availability

Raw data, and the Python code used for image analysis can be accessed at Zenodo repository (DOI:https://doi.org/10.5281/zenodo.8095546). All other data and analyses are included in the main text or in SI Appendix.

## ACKNOWLEDGMENTS

IB, NK, BMF were supported by the Deutsche Forschungsgemeinschaft (DFG, German Research Foundation) under Germany’s Excellence Strategy - EXC-2068-390729961. BMF acknowledges support through a Heisenberg grant of the DFG (FR3429/4-1,4-2). We thank Carl Modes for stimulating discussions as well as Szabolcs Horváth for advice on image analysis.

## Supporting Information for

## This PDF file includes

Supporting text

Figures S1 to S10

Tables S1 to S3

Legends for Movie S1

## Other supporting materials for this manuscript include the following

Movie S1

Dataset S1

Software S1

## Supplementary text

### Image analysis of TEM data

#### Preprocessing

A total of 38 TEM images of nascent valves were obtained by extracting valve SDVs from synchronized cell cultures as described previously [1]. Of these, 23 TEM images required manual preprocessing. This included changing the image contrast and sharpness in the GIMP image redactor using linear rescaling of intensity values. The GIMP image redactor was also used to manually remove impurities. Fifteen TEM images did not require any manual preprocessing and were used directly for skeleton recognition. A typical TEM image is shown in Fig. S1A.

#### Skeletonization

Next, TEM images were analyzed using the python scikit-image library (v0.15.0) [2]. To speed up analysis, images were re-scaled to 80% of their original size. In a first step, images were smoothed using a median filter with diameter of 40 nm, followed by adaptive thresholding with a window size of 155 nm. This resulted in a binary mask that included the contour of the silica valve and occasional small objects of relatively high electron density (Fig. S1B). Correspondingly, small objects with a total area smaller than 0.75 μm^2^ were removed from the mask. This curated mask was then skeletonized using Zhang’s method (Fig. S1C) [3]. From this skeleton, short side-branches with length less than 50 nm length were removed using a custom designed skeleton pruning algorithm. Small cycles enclosing an area of less than 490 nm^2^ (presumably corresponding to cribrum pores) were filled in, and the resultant binary mask was skeletonized again. In this way, we ensured that spurious branches encasing cribrum pores would not be considered in the further analysis. As a result, we obtained a pruned skeleton, devoid of short branches and most cycles (Fig. S1D). In total, we were able to skeletonize 38 valve SDVs (Fig. S2).

#### Rib spacing quantification

The spacing between neighboring ribs was calculated from skeletonized valve networks as follows. First, the original skeleton was dilated with a disk element of radius 2 pixels. This dilated mask was then inverted, and the median-axes function applied. This provided a mask of pixels that are equally distant to adjacent ribs, together with the scalar value of this equal distance *d/2* for each pixel (Fig. S3).

#### Topological analysis

Topological analysis of valve skeletons was implemented using the mahotas python library (v1.4.11) [4]. Specifically, we classified pixels of the skeleton according to pixel connectivity: pixels with 1-pixel connectivity were categorized as tip points of ribs, pixels with 3-pixel connectivity were classified as branch points (Fig. S4A). This provided the total number of rib tips and branch points.

Each pixel of the skeleton was then categorized, thus obtaining the number of tips and branching points for the entire skeleton. Transverse cross-connection between radial ribs were automatically identified as skeleton paths connecting two branching points when their paths were shorter than 170 nm and they exhibited an 60° to 90° angle relative to the local radial direction (Fig. S4B, green lines). To restrict the analysis to branch points of radial ribs, any branching points connected to transverse cross-connections as well as branching points inside the central valve area (25% of the total valve radius) were excluded from further quantitative analysis. The valve center point was calculated as center mass point of the convex hull of the skeleton of the rib network.

### Numerical methods

#### Numerical integration of partial differential equations

To numerically integrate the model given in Eqs. [1-4], we used a forward Euler scheme with constant time step and regular grid, with three-point stencil for the diffusion operator and reflecting boundary conditions. Integration was carried out in a large static square domain (Figs. 2 and 3), or in a dynamic disk-shaped domain that expands at constant speed (Fig. S7). To attenuate grid effects of the square grid used in simulations, we added a noise term in the diffusion coefficients of all components as described in [5,6]. Specifically, we defined diffusion coefficients at each position and at each time step as follows: *D(x,y)=D*_*i*_*(1+Δ(x,y))*, where *Δ(x,y)* denote independent, random variables uniformly distributed in interval of *(−1,+1)*. Values of all model parameters in the simulation of valve patterns of *T. pseudonana* are listed in Dataset S1 as excel spreadsheet.

#### Initial conditions

For simulations of valve morphogenesis in *T. pseudonana* the PSS was digitally drawn by hand and then imported into simulations as initial condition. The PSS size was re-scaled to match the average effective diameter of the analyzed PSS obtained from the TEM images. We confirmed that similar patterns emerge with the same number of initial rib stems when a perfect annulus of same width is chosen as initial condition instead of a distorted annulus (Fig. S5). We further confirmed in preliminary simulations that adding small-amplitude noise to the initial concentration of the intermediate component does not change results (Fig. S10).

For the simulations of valve patterns in the pennate diatom *A. sibiricum* model as shown in Fig. 4B in the main text, we used an elongated seed of two long ribs separated by narrow slit as initial condition. This PSS featured a corrugated, rough surface (Fig. S9), which had been simulated according to a random model: the height profile *y*(*x*) of the PSS surface (with PSS aligned along the x-axis) was represented as a random Fourier series 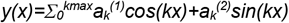 where 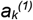 and 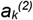 are independent normally distributed random numbers with standard deviation *w* and the sum over wavevector *k* running from 0 to *k*_max_ in increments of *k*_max_ *L/π*. The model parameters used in the simulations of valve patterns of *A. sibiricum* are listed in Dataset S1 as excel spreadsheet.

For the simulations of valve patterns of the centric diatom *C. cryptica* shown in Fig. 4B in the main text, initial conditions were chosen as a thin annulus analogous to the simulations for *T. pseudonana*. After the initial thick stems had formed, the values of model parameters were changed to new values that produce thin ribs.

On a technical note, this switch of parameters was realized by running two simulations subsequently, where the spatial concentration fields of all components at the end of the first simulation were used as initial condition for the second simulation. The model parameters used for the simulation of valve patterns of *C. cryptica* model are listed in Dataset S1 as excel spreadsheet.

#### Expanding SDV domains

To address the impact of dynamic spatial confinement as represented by the expanding SDV, we performed additional simulations in a growing domain with reflecting (i.e., no-flux) boundary conditions. The dynamics of this growing domain is characterized by additional parameters: the initial offset of the SDV boundary with respect to the contour line of the PSS, the relative speed of expansion of the SDV relative to the speed of rib pattern propagation, and a non-zero degradation rate *k*_*i*_ of inhibitor. We investigated four scenarios: branching dynamics with/without inhibitor degradation and with/without expanding SDV boundary (Fig. S7).

For these simulations, the propagation speed of silica pattern in an unbounded domain was determined in an initial step as follows: using simulations analogous to those shown in Fig. 2B of the main text, we tracked the mean radial position of the isoline of inhibitor concentration at *[I]=0*.*25* as function of time. For the simulations in an expanding domain shown in Fig. S7, we set the speed of SDV expansion equal to this pattern propagation speed. The initial distance between the SDV boundary and the PSS pattern was set to 150 nm for simulations. The values of the kinetic parameters are stated in the figure caption of Fig. S7.

## Supplementary figures

**Figure S1.**
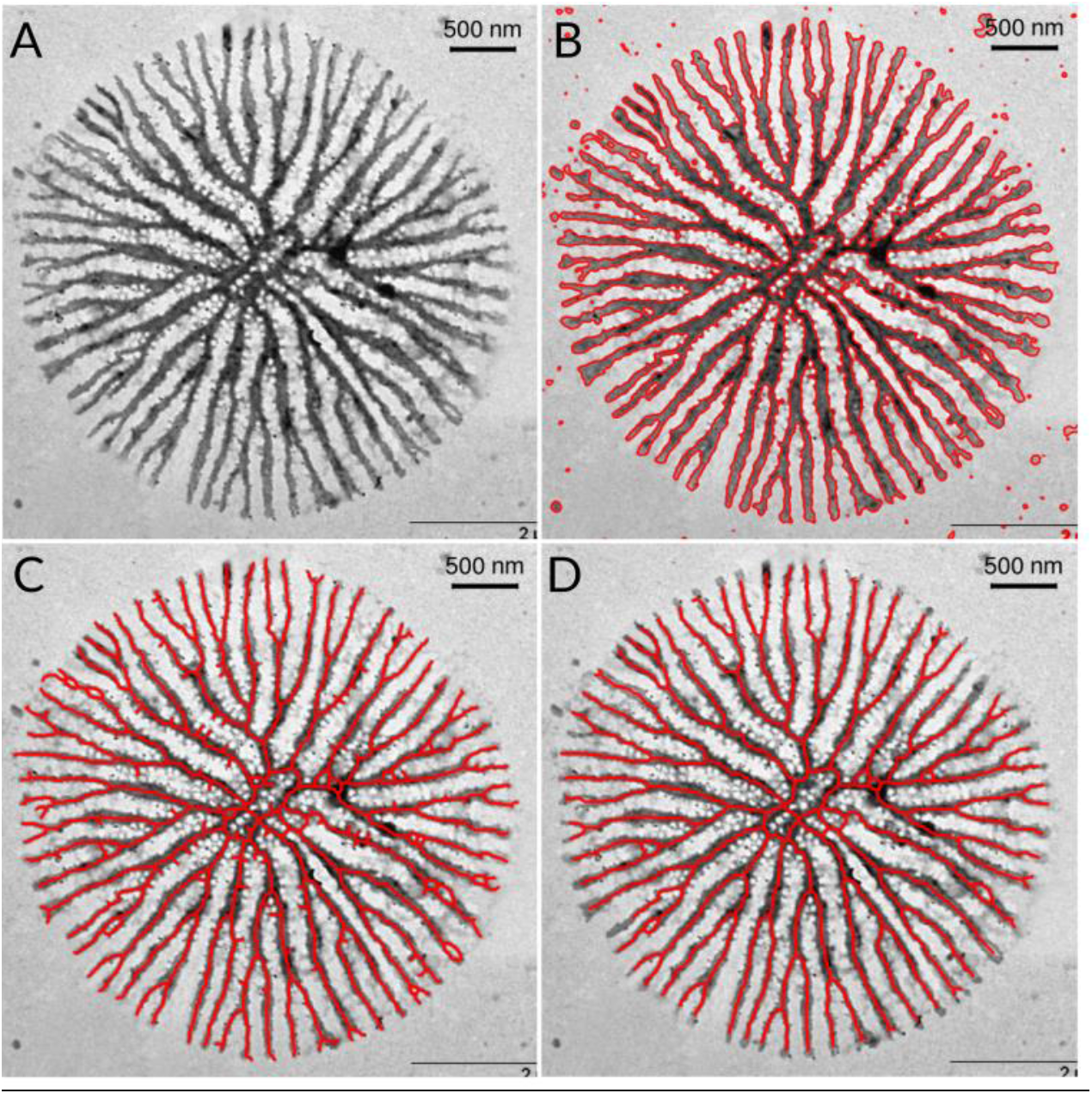
Analysis of TEM images. (A) Preprocessed image with removed impurities. (B) Binary mask obtained by adaptive thresholding (red). (C) Skeletonization of the binary mask from panel B. (D) Skeleton from panel C after pruning of short branches and removal of small cycles.

**Figure S2.**
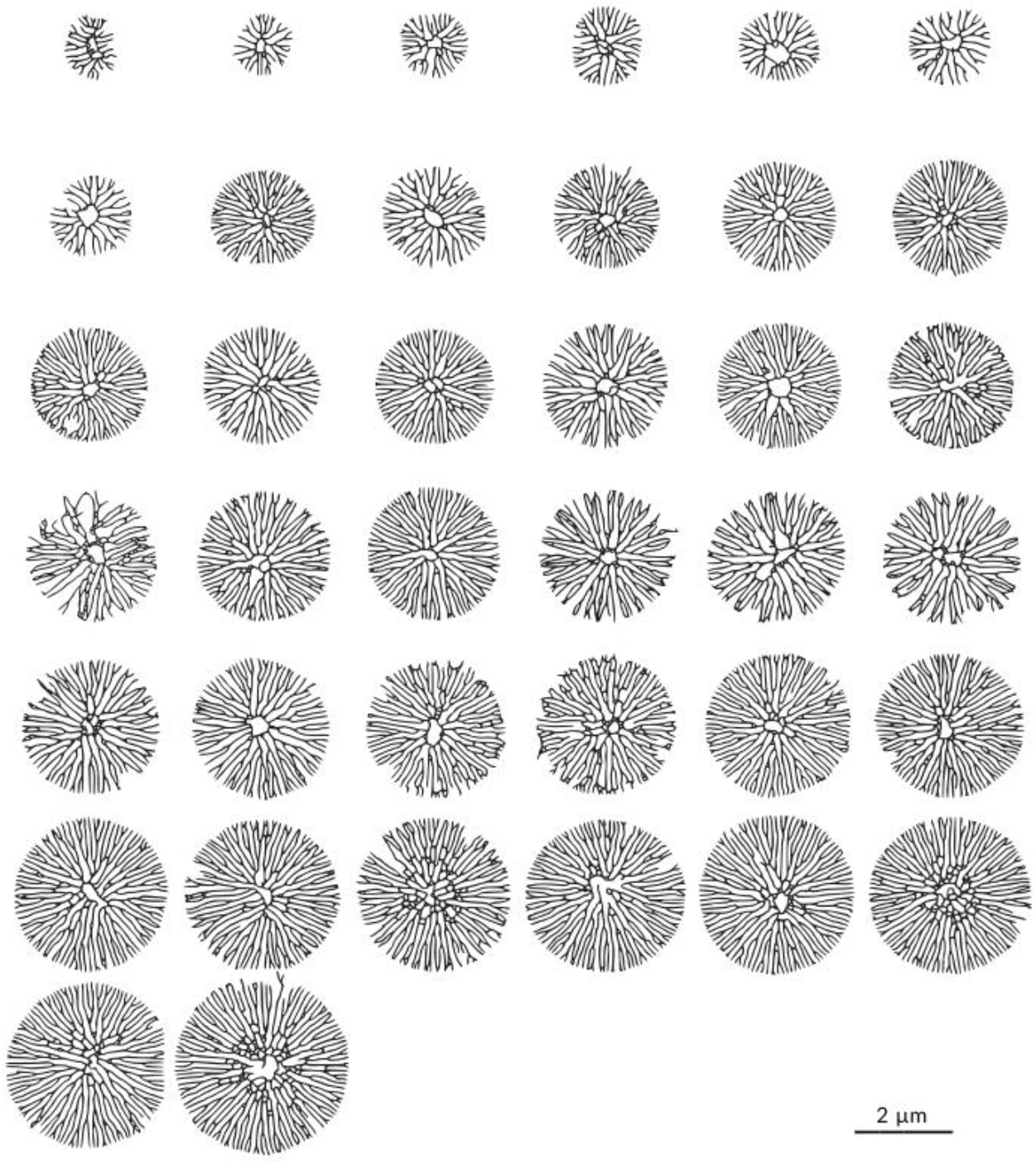
Valve skeletons. The images show the silica skeletons obtained from TEM micrographs of *T. pseudonana* valve SDVs that were used to obtain morphometric data as shown in Fig. 2 of the main text. The skeletons were sorted according to ascending valve radius.

**Figure S3.**
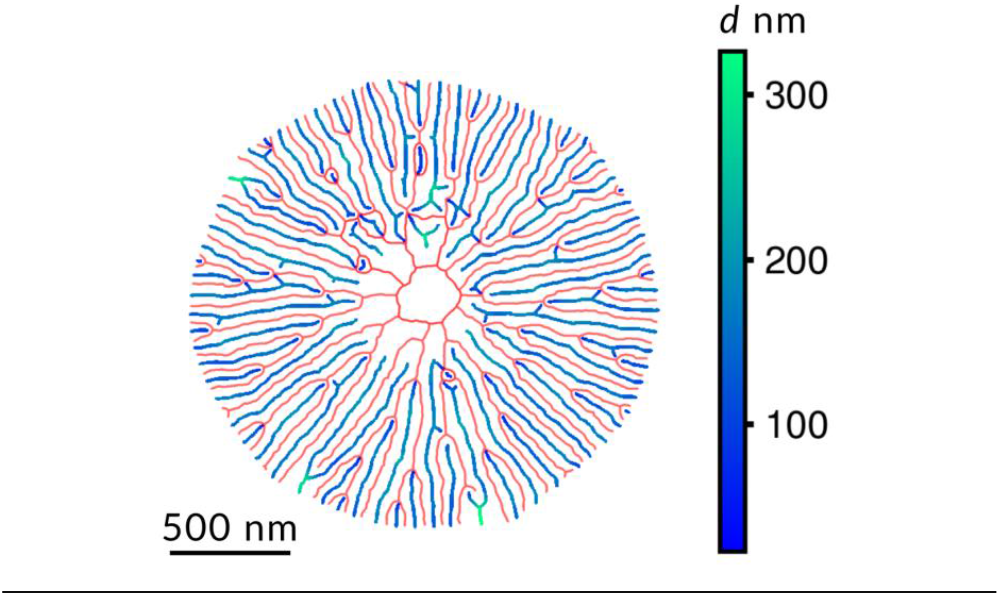
Local inter-rib spacing of a typical valve skeleton. The original valve skeleton is shown in red. The graph in blue-green represents all pixels with equal distance *d*/2 to adjacent ribs as obtained by the median-axes function. The blue-green color code indicates this distance. We used *d* as proxy for the local inter-rib spacing as reported in Fig. 2C in the main text.

**Figure S4.**
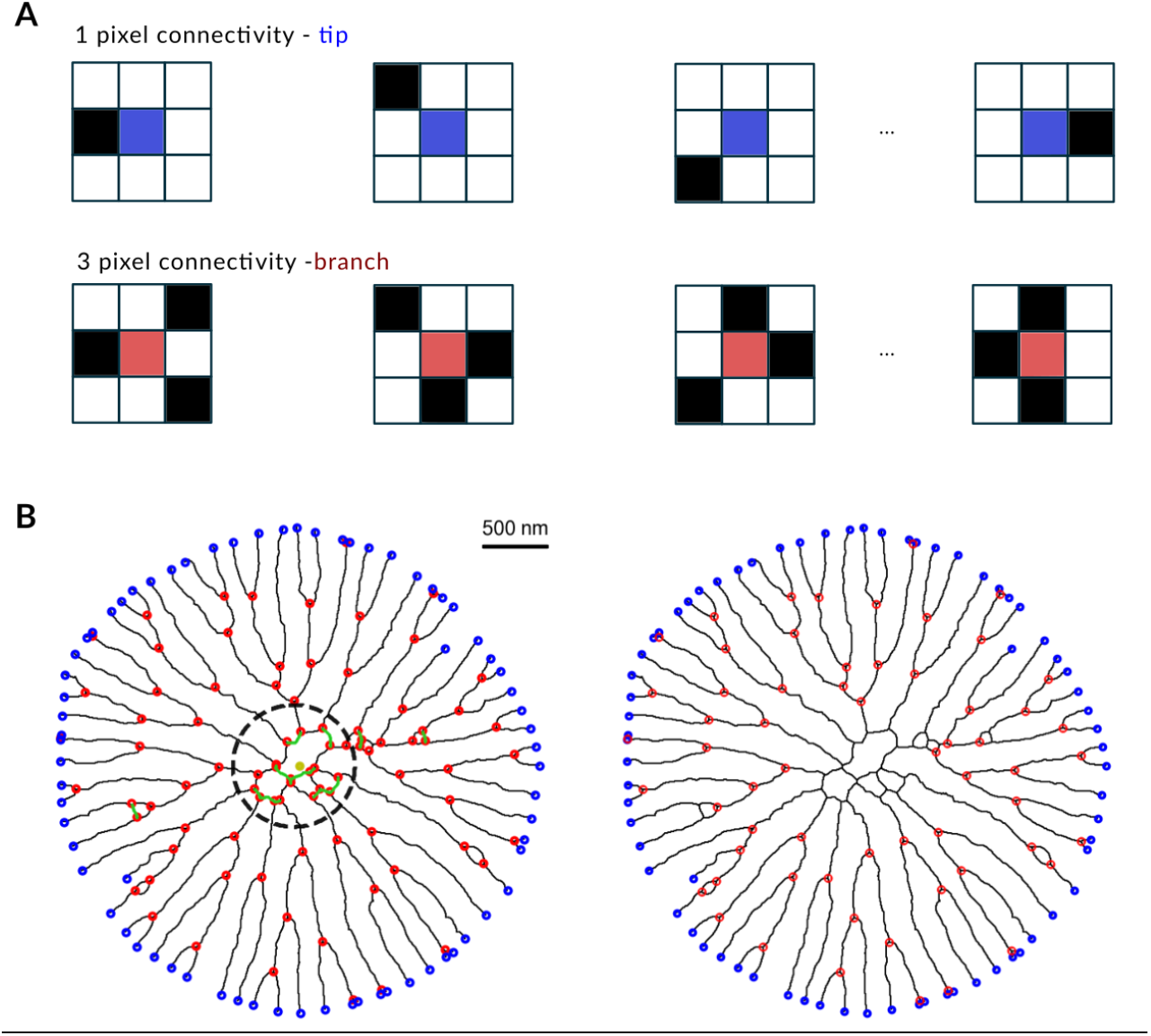
Topological analysis of valve skeletons. (A) *Top:* Representative 2D-kernels used to detect pixels marking tip points of skeletonized ribs by pixel-connectivity analysis. *Bottom:* Representative 2D-kernels used to detect pixels marking branch points. (B) *Left:* Example classification of pixels for a typical valve skeleton highlighting tip points (blue), branching points (red). Transverse cross-connections between radial ribs were determined as skeleton paths connecting two branching points of length shorter than 170 nm and angle between 60° and 90° relative to the local radial direction (green). Branching points inside the area enclosed by the dashed circle (radius equal to 25% of effective valve radius) as well as branch points connected to transverse connections were not included in subsequent quantifications. *Right:* Tip points (blue) and branch points (red) used for quantification as in Fig. 2 in the main text.

**Figure S5.**
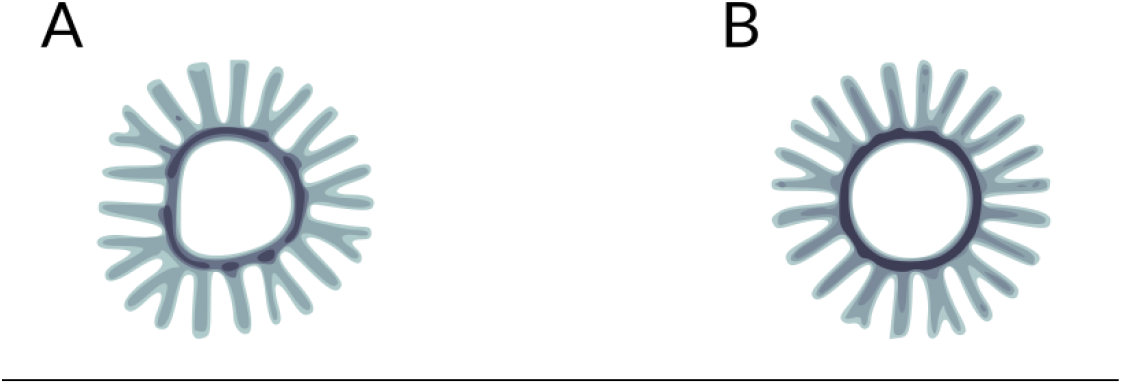
Number of emerging rib stems for different initial conditions. (A) Early stage of a simulated rib pattern for a digitally-drawn distorted annulus as initial condition as used in Fig. 2B in the main text, resulting in n=16 rib stems. (B) Same as in panel A, but using an ideal circle of same width as initial condition. The same number of rib stems are produced as in A.

**Figure S6.**
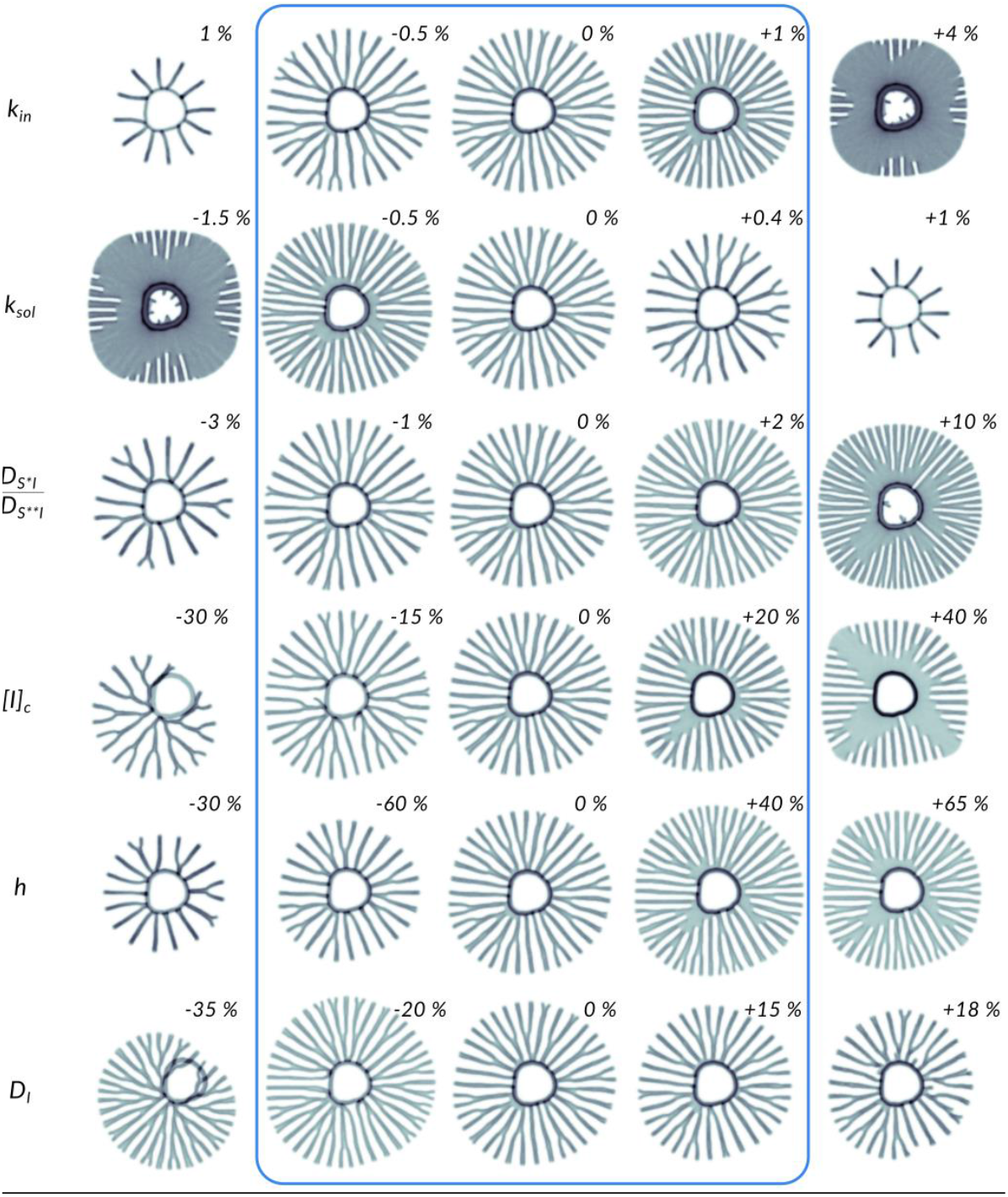
Sensitivity analysis of model parameters. Each row displays simulation results for the minimal model of rib pattern morphogenesis with a single parameter changed as indicated. The relative change of the corresponding parameter in percent is shown next to each simulated rib pattern. The blue box indicates a region of parametric robustness, where patterns change only insignificantly. The data show that the parameters of the Gray-Scott model describing conversion of precursor to intermediate must be tuned precisely to a regime known to produce a self-replicating dot pattern. Yet, the model is robust with respect to parameters characterizing the inhibitor feedback, which represents the essential and novel ingredient of our model.

**Figure S7.**
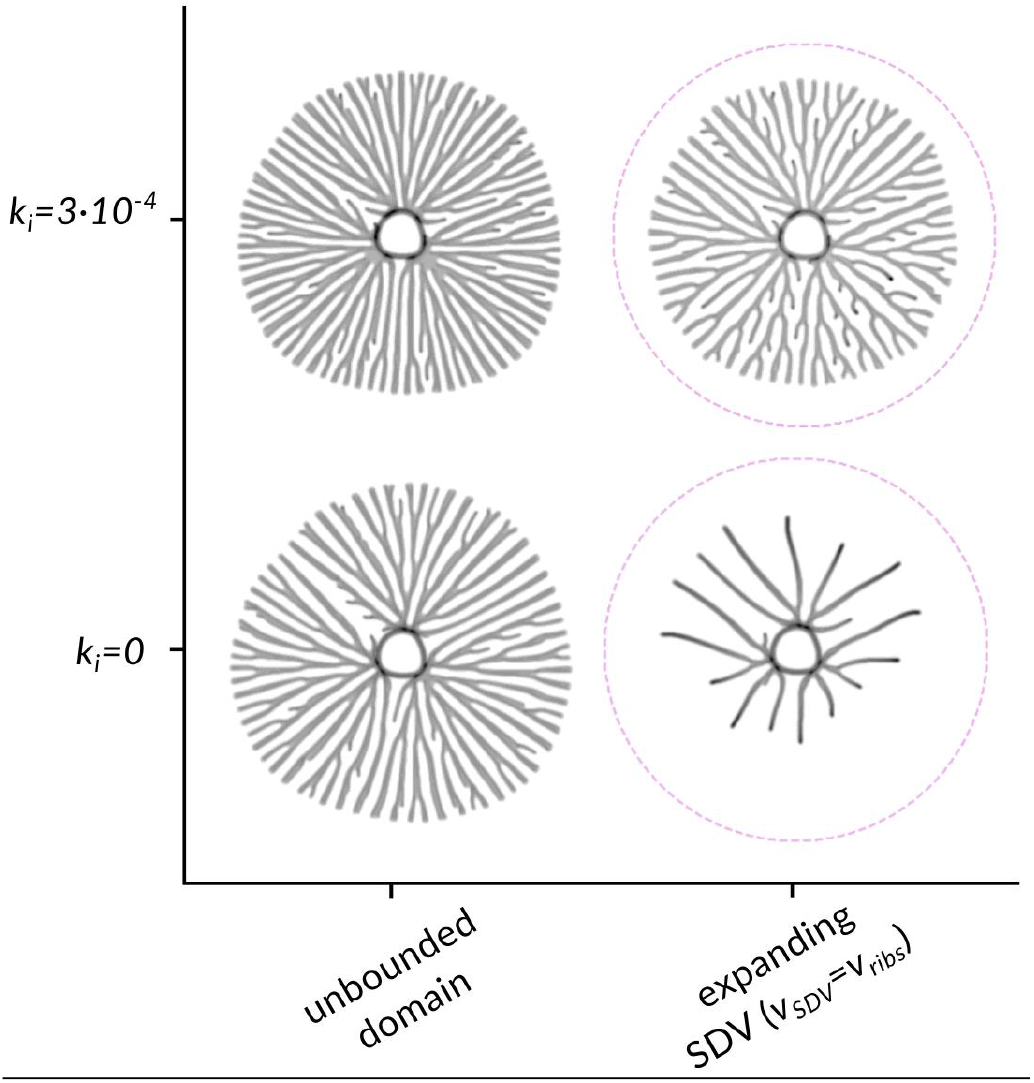
Simulated rib patterns in unbounded and expanding domain, with and without inhibitor degradation. Simulated dynamics of silica branching morphogenesis in an effectively unbounded domain and in an expanding domain with reflecting boundary conditions. For each boundary condition, we consider a scenario without inhibitor degradation (*k*_*i*_=0) and with a non-zero degradation rate of inhibitor (*k*_*i*_*>0*). Parameters: *k*_*in*_ *= k*_*out*_ *= 0*.*0073, k*_*sol*_ *= 0*.*135, I*_*c*_ *= 5*.*5, h = 50, D*\* *= 0*.*112, D*\*\**= 0*.*0552, D*_*i*_ *= 0*.*17*.

**Figure S8.**
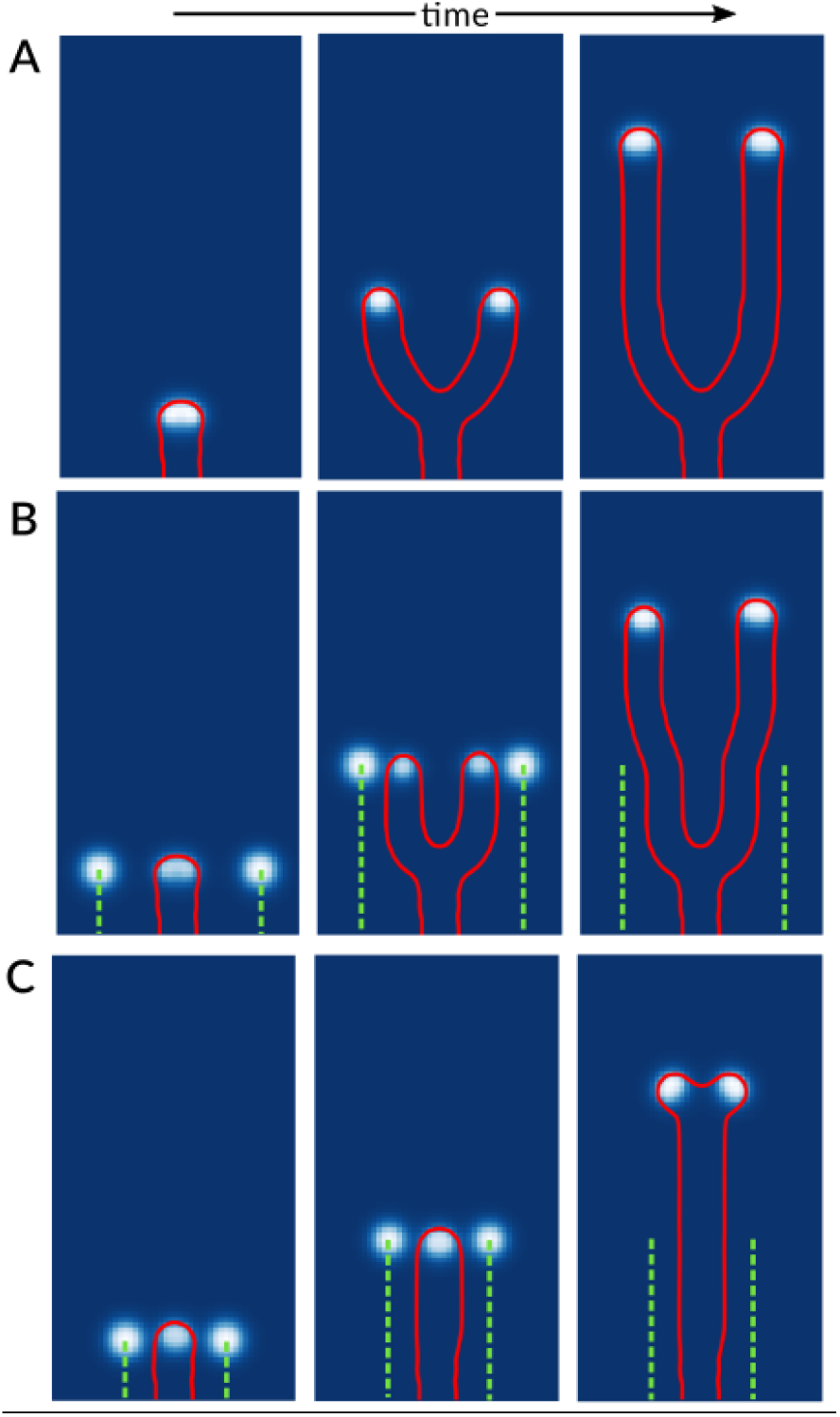
Normal rib branching in the presence of artificial precursor sinks. (A) Normal rib propagation and branching. This precursor concentration [*S*^*^*I*] is shown in blue color code with isoconcentration curve in red. (B) Same as panel A, but with artificially imposed “ghost ribs” consisting of sinks of precursor that propagate at the same speed as normal rib propagation. The simulated central rib branches despite the presence of the precursor sinks. Once precursor sinks are removed, the shape of the branch relaxes to that of a normal branch without precursor sink. (C) Same as panel B, but with distance of “ghost ribs” equal to 60% of the normal rib-rib spacing. In this (unachievable) situation, branching of the propagating central rib is suppressed and only resumed once the precursor sinks are removed.

**Figure S9.**
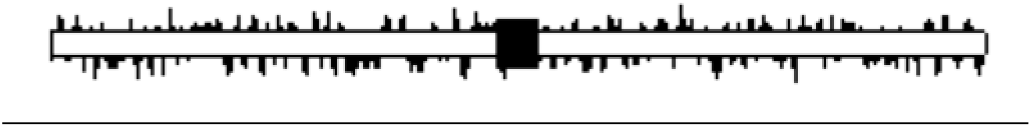
PSS configuration used for the pennate diatom *A. sibiricum*. Concentration pattern of the intermediate S^**^·I that was used as initial condition for the simulations of valve morphogenesis in the pennate diatom *A. sibiricum* as shown in Fig. 4B in the main text.

**Figure S10.**
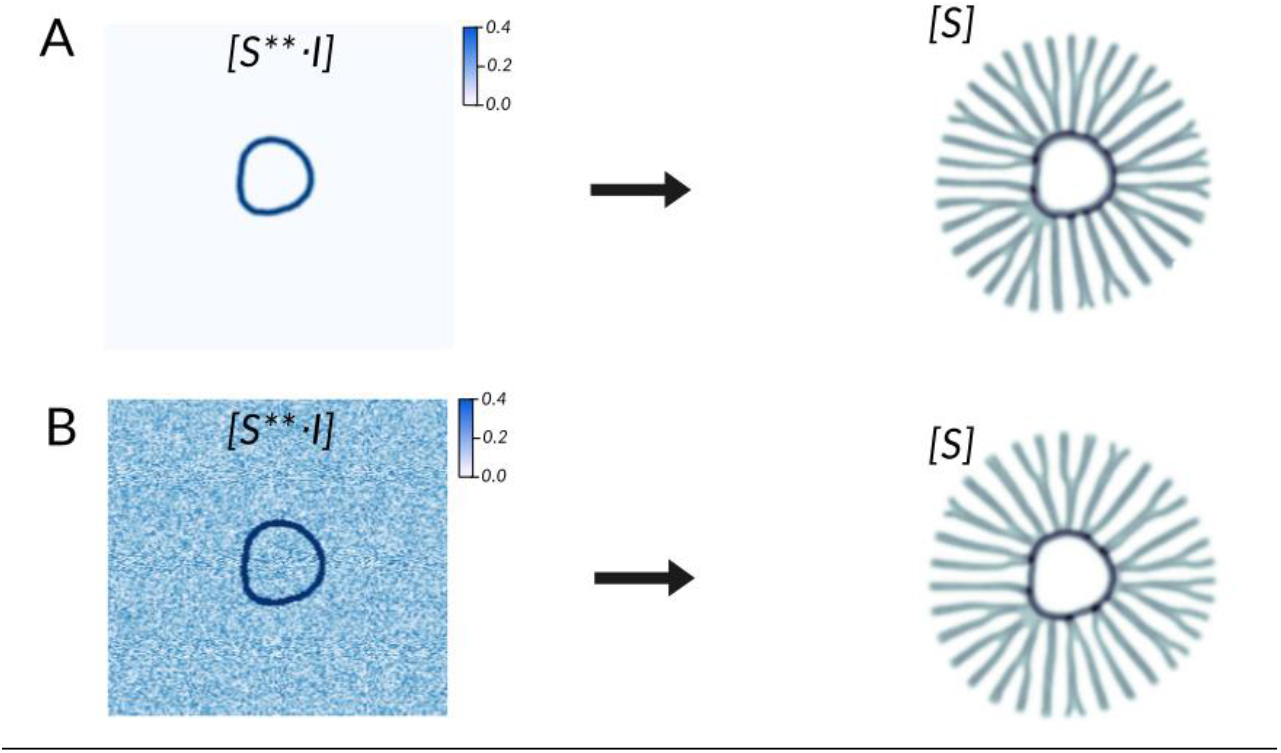
Robustness to concentration fluctuations. (A) *Left:* Initial seed consisting of a distorted annulus of elevated concentration of intermediate *S*^**^·*I* as used for simulations in Fig. 2B of the main text. *Right:* Corresponding simulated rib pattern consisting of solid silica. (B) *Left:* Same as panel A, but with random concentration fluctuation added (independent random numbers uniformly distributed in the interval [0,0.3] added to each grid point). *Right:* the resultant simulated rib pattern shows no significant difference from the pattern without noise shown in panel A.

## SI legends

**Data set S1 consisting of:**

**Table S1 (excel spreadsheet)**. Morphological quantification of TEM images of valve SDVs

**Table S2 (excel spreadsheet)**. Morphological quantification of simulated silica patterns.

**Table S3 (excel spreadsheet)**. List of model parameters used in simulations.

**Movie S1**. Simulation of silica valve morphogenesis showing the normalized concentration of solid silica as function of time.

